# Small language models enable rapid and accurate extraction of structured data from unstructured text: an example with plants and their specialized metabolites

**DOI:** 10.1101/2025.05.19.654923

**Authors:** Lucas Busta, Alan R. Oyler

## Abstract

Transformer-based large language models are receiving considerable attention because of their ability to analyze scientific literature. Small language models (SLMs), however, also have potential in this area, have smaller compute footprints, and allow users to keep data in-house. Here, we quantitatively evaluate the ability of SLMs to: (i) score references according to project-specific relevance and (ii) extract and structuring data from unstructured sources (scientific abstracts). By comparing SLMs’ outputs against those of a human on hundreds of abstracts, we found that (i) SLMs can effectively filter literature and extract structured information relatively accurately (error rates as low as 10%), but not with perfect yield (as low as 50% in some cases), (ii) that there are tradeoffs between accuracy, model size, and computing requirements, and (iii) that clearly written abstracts are needed to support accurate data extraction. We recommend advanced prompt engineering techniques, full-text resources, and model distillation as future directions.

## 1. Introduction

Language models are emerging as powerful tools for a wide array of tasks, with a particularly promising role in processing scientific literature (Agathokleous et al. 2024; Jin et al. 2024; Lam et al. 2024; Simon et al. 2024; Busta et al. 2024b; Knapp et al. 2024a). Scientific articles compile results from decades, if not centuries, of effort by scientists worldwide. However, the automation of classification, summarization, and data extraction tasks related to this literature remains a challenge because natural language is a complex data type. In other fields with intricate data, such as image and sound, a proven strategy is to build mathematical models of the input data type that can then be leveraged to summarize, classify, or otherwise manipulate the input. Modeling natural language is a long-standing field of study, but recently, the development and increase in accessibility of transformer-based language models have led to substantial advances in our language processing ability. Perhaps we can solve some of the many challenges with automated processing of scientific literature by applying transformer-based language models.

A considerable number of recent investigations are focused on applying large language models to scientific literature (Jin et al. 2024; Busta et al. 2024b; Shiu and Lehti-Shiu 2024; Sarumi and Heider 2024; Knapp et al. 2024a). For example, large language models have been utilized to perform tasks such as text classification, text summarization, and question answering (Dalal et al. 2024; Riordan 2024; Shiu and Lehti-Shiu 2024; Guo et al. 2023; Yin et al. 2019). Generally, these large models require significant memory—hundreds of gigabytes—to store high billions or trillions of parameters required at runtime. However, a diverse range of language models exists beyond the popular large models from, for example, OpenAI, Anthropic, Google, and Mistral. In particular, small language models (SLMs) have gained attention due to their smaller sizes (low billions or even just millions of parameters) and thus reduced computing requirements. Furthermore, though the small models are not as general purpose as the large models, the emerging evidence suggesting the small models are effective in various, albeit specific natural language processing tasks (Lepagnol et al. 2024; Guo et al. 2023; Zhu 2024; Lewis et al. 2019). Thus, these small language models are intriguing because they suggest that individual scientists could use them on ordinary personal computing devices to potentially enhance scientific literature processing tasks. Importantly, running the small models on local hardware also avoids passing private and/or copyrighted content to large language model companies, which is prohibited by many research institutions and industrial organizations.

In the present work, we aimed to develop and evaluate a proof-of-concept small language model processes to support the expansion of databases that document plants and the specialized metabolites that each may produce. Other databases have been created in the past to document this same type of information (Zeng et al. 2024; Tay et al. 2023; Gallo et al. 2023; Rutz et al. 2022; Sorokina and Steinbeck 2020; Nguyen-Vo et al. 2020; Yang et al. 2019; Chen et al. 2017; Xie et al. 2015), but these databases, so far, do not leverage the potential provided by language models. We experimented with models to conduct two major tasks: (i) scoring articles based on their relevance to need-specific criteria (in this case, whether they contained reports of a specific plant making a specific chemical compound) and (ii) extracting and structuring information on the occurrence of specific chemical compounds in specific plant species. We tested a dozen language models’ abilities on these tasks by manually reading, labelling, and extracting data from more than 100 to more than 1000 scientific abstracts, depending on the task, then measured the models’ ability to perform those same tasks. Overall, our findings indicate that small language models, while not perfect, effectively aid in filtering scientific literature references and in extracting data. We recommend that researchers both experiment with these models and monitor for updates in literature processing software that incorporate language model-enabled features.

## 2. Results and Discussion

To develop and evaluate a potential role for small language models in creating a phytochemical occurrence database, we assessed such models’ abilities with regard to two tasks: (i) to quickly score references according to whether the reference reports the occurrence of a specific compound in a specific plant species (Task 1, Section 2.1), and (ii) to evaluate language models’ ability to extract an experimentally-supported compound occurrence dataset (Task 2, Section 2.2). For these investigations, we chose to use six triterpenoid compounds as test cases (**Fig. 1A**). The six triterpenoid test cases under study here presented a challenge because they have been mentioned in the literature (going back to the 1960s) by many names. Indeed, CAS SciFinder® indicates that a total of more than 52 names has been associated with these six compounds, potentially complicating efforts to retrieve references describing the occurrence of specific plant chemicals. Fortunately, triterpenoids (and the vast majority of all other chemical entities) are identified explicitly by their CAS Registry® numbers (**Fig. 1A**), which means that references to a given compound that use varied nomenclature can be collected simultaneously and non-ambiguously when using CAS Registry® number-based search strategies. While other identification number systems exist, such as PubChem® and LOTUS numbers, these alternate systems are not as comprehensive as CAS Registry® numbers. Thus, where possible, searches with identification numbers, as opposed to common names, are preferred because this approach ensures not only that a broader array of references is retrieved, but also that those reference relate to one and the same compound, including the correct stereochemistry.

**Figure 1.**
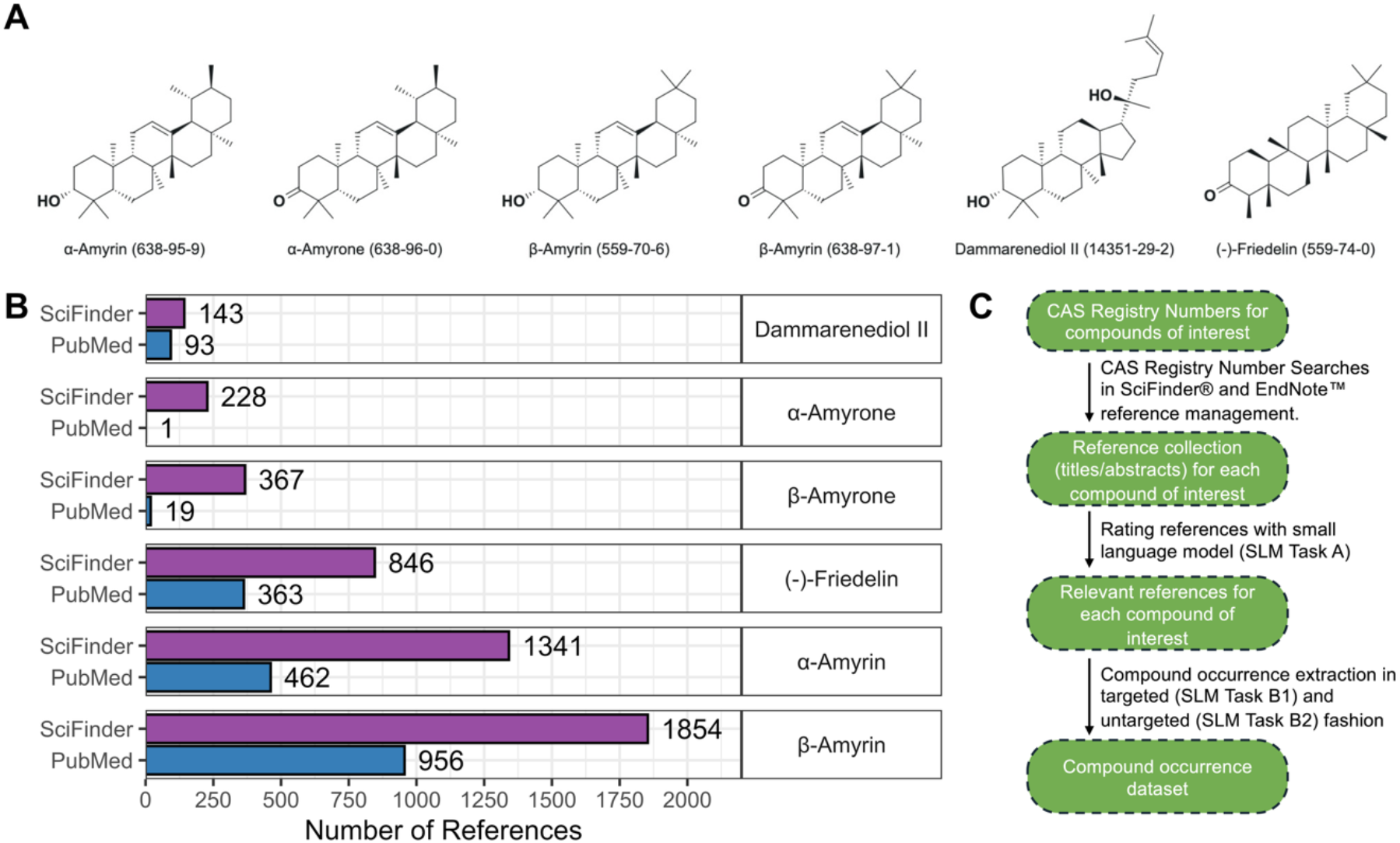
Comparison of SciFinder® versus PubMed® as a data source and schematic of the small language model workflow for retrieving compound-species associations from literature. **A**. Structures, common names, and CAS Registry® Numbers for the six triterpenoid compounds used as test cases in our small language model development and evaluation work. **B**. Bar plot comparing the number of references (x-axis) found by SciFinder® and PubMed® (y-axis) for the six different triterpenoids (vertically arranged panels) studied in this work. Each bar represents the number of references found by the indicated search tool for a particular triterpenoid. The absolute number of references found is shown in text to the right of each bar. Bars are color coded according to search tool (SciFinder® in purple and PubMed® in blue). SciFinder® searches were conducted using CAS Registry® Numbers, while PubMed® (which does not generally use these registry numbers) searches were conducted using compound common names. **C**. Schematic for the workflow we developed to extract compound occurrence data from information in the literature. Files or information are shown in green bubbles, while steps or actions are shown as arrows. The workflow consists of searching the literature with SciFinder® based on CAS Registry® numbers then creating a repository of references and associated full text PDF files in an EndNote^TM^ database; then filtering references for those of highest task-specific relevance (SLM Task A) and finally extracting compound occurrence data in either a targeted (SLM Task B1) or untargeted (SLM Task B2) fashion. Abbreviations: SLM: small language model.

To obtain references describing our six triterpenoids of interest, we used CAS Registry® numbers to search SciFinder®, which, although requiring a subscription, allows the user to enter a CAS Registry® number and then navigate directly to literature references that relate to that specified compound. PubMed®, although providing open access, does not generally support searches based upon CAS Registry® numbers or PubChem ID numbers, so we conducted searches in PubMed using compound common names. We first considered the two most common compounds in our case study set, α and β-amyrin. In SciFinder®, we found over 1,340 and more than 1,850 hits for these two compounds, respectively, compared to fewer than 500 and 1,000 hits in PubMed® (**Fig. 1B**). Results were similar for the other four triterpenoid test cases (**Fig. 1B**). In total, ∼3,200 SciFinder® references were retrieved using our searches, while ∼1,500 references were retrieved by PubMed®. Therefore, we used SciFinder®-retrieved references to develop and evaluate small language model-based reference ranking and occurrence dataset extraction processes (**Fig. 1C**).

### 2.1 SLM Task A: Rating references according to relevance with a small language model

At this stage in the present work, we had used SciFinder® to collect more than 3,000 references associated with one or more of the six triterpenoids that comprised our test cases for compound occurrence data collection. Our first aim was to determine the efficacy of small language models with respect to filtering the references for articles of interest. In this case, our interest was in articles that reported phytochemical occurrences (i.e., evidence for a specific plant species producing a specific chemical compound). To establish a benchmark against which to evaluate small language model performance we read more than 1,500 of the references in our collection, including their titles and abstracts, and classified each as “reporting an occurrence”, “maybe reporting an occurrence”, or “not reporting an occurrence” (**Supplemental File 1**). These human-read citations included all the reference citations for α-amyrone, β-amyrone, dammarenediol II, as well as (−)-friedelin. For an article to be considered as “reporting an occurrence” its title or abstract needed to indicate that the article in question provided experimental evidence for the presence of a particular plant chemical in a particular plant species. Articles whose titles or abstracts merely contained co-occurrences of a plant chemical name and a plant species name without indicating that there was experimental evidence for an association between the two were classified as “not reporting an occurrence”. Citations that did not explicitly indicate that their articles contained experimental evidence for a compound’s occurrence but instead implied that such evidence might be present in the full text (to which we did not have access) were classified as “maybe reporting an occurrence”. Of the 1,558 references that we read, 720 were classified as “reporting an occurrence” (46%), 332 were classified as “maybe reporting an occurrence”, (21%) and 506 were classified as “not reporting an occurrence” (33%).

We next evaluated how well language models could classify references according to whether they reported the occurrence of a phytochemical using the 1,558 manually classified references as a ground-truth set. We used the bart-large-mnli model, selected because it is one of the most downloaded on Huggingface.co, a major hub for open-source language development, largely due to its versatility and high speed – we found that it could process 45,000 articles / hour, a desirable characteristic for a model that will be used to filter inputs into a multi-step processing pipeline. This small language model is employed by providing it with a body of text and then one or more classifier phrases. The model then assigns a score to each phrase to indicate how closely that phrase relates to the provided text. The bart-large-mnli model card (i.e., the instruction manual) suggests presenting the model with a classifier phrase framed as a hypothesis (e.g., “This text is about politics”). Accordingly, we investigated phrases such as “Amyrin is present in plants” as well as paired phrases in which a hypothesis was matched with the exact negative (i.e., “Amyrin is present in plants” and “Amyrin is not present in plants”). Our early experiments showed that composite scores derived from the pairs’ individual scores improve the signal-to-noise ratio in the classification task. Furthermore, we noted that multiple compound names could be included in these positive and negative phrases (for instance, “friedelin, friedooleanan-3-one, friedelan-3-one, friedelanone, or friedeline is found in plants”; full classifier phrase details are provided in **Supplemental File 2**). In future large-scale operations, a single, general classifier phrase, which is not based on compound names, would be preferred if the performance was comparable to that of our specific classifier phrase system, which is based on compound names. Therefore, we also tested the more general classifier phrase, “The text discusses plants that contain specific compounds.”

Using the two classifier phrase approaches described in the previous paragraph, we instructed the bart-large-mnli model to assign two scores to each of the 1,558 references, one composite score from the binary/two classifier phrase system, as well as a score for the general classifier phrase. Composite scores (means and standard deviations) for, respectively, references that reported occurrences / maybe reported occurrences / did not report occurrences for (−)-friedelin were 0.9 ± 0.1, 0.8 ± 0.1, and 0.7 ± 0.1 (**Fig. 2A**, top panel). Results were similar for the other three triterpenoids (**Fig. 2A**). Scores from the general classifier phrase for, respectively, references that reported occurrences / maybe reported occurrences / did not report occurrences for (−)-friedelin were 0.9 ± 0.06, 0.9 ± 0.04, and 0.8 ± 0.2 (**Fig. 2B**, top panel), and again, results were similar for the other three triterpenoids (**Fig. 2B**). This illustrates that the two-classifier phrase system and the general classifier phrase system both worked comparably well among references describing four different triterpenoid compounds and may also to a similar extent for compounds other than triterpenoids.

**Figure 2:**
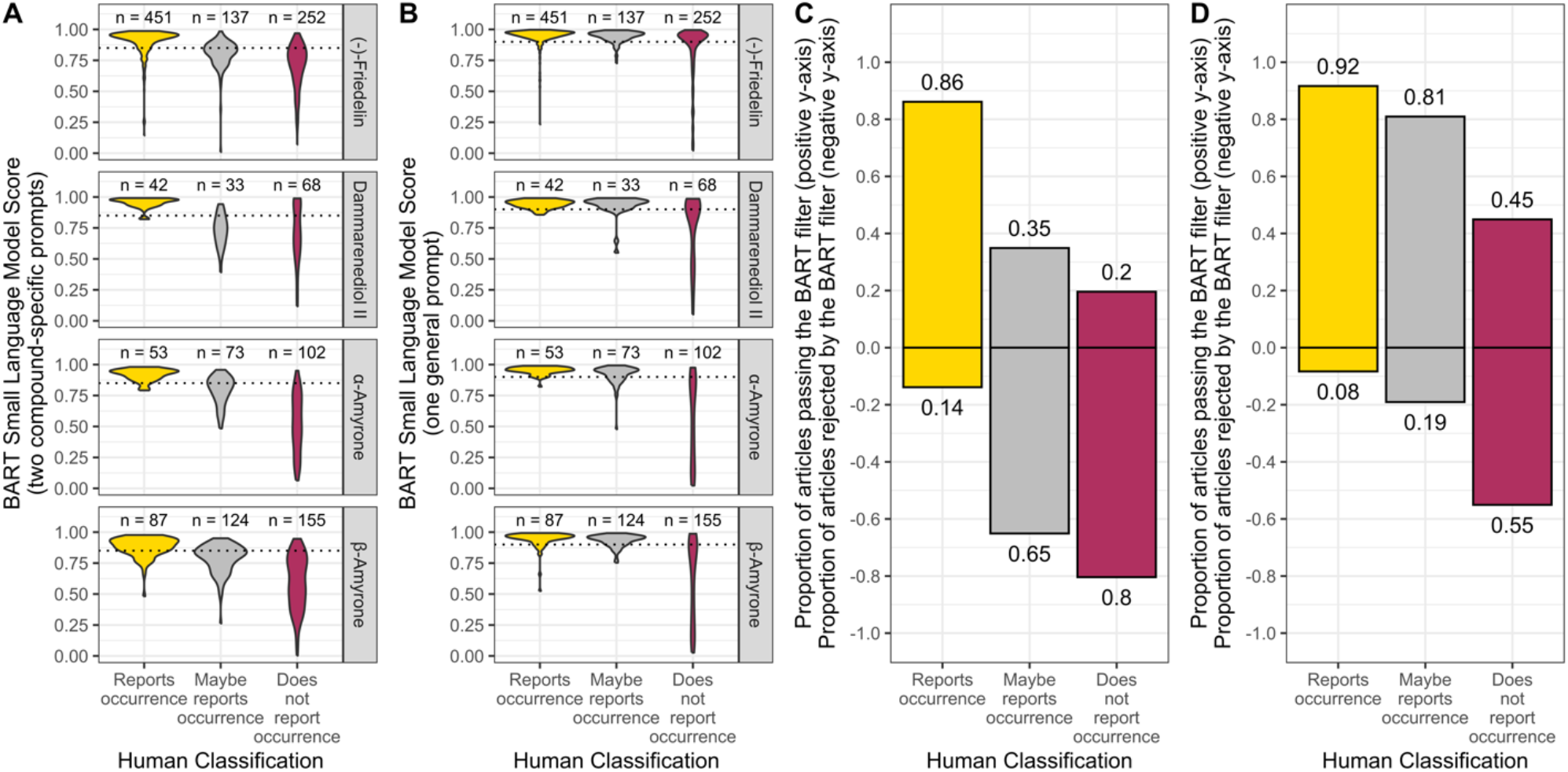
Performance of small language models on a reference relevance ranking task. **A and B**. Violin plot showing the score (BART Small Model Language Score, y-axis) assigned to references by the bart-large-mnli small language model. Scores range from zero (low relevance) to one (high relevance) and indicate the relevance of a given reference to a user-defined natural language criterion. In panel A, the score is derived from two, chemical compound-specific criteria (full details in methods section), while in panel B, the score is derived from a single, generic criterion (“chemical compounds are found in plants”). In both panels, scores are broken out according to whether the reference was labeled by a human as “reporting an occurrence”, “maybe reporting an occurrence”, “not reporting an occurrence” of a specific chemical compound in a specific species (x-axis). The number of references belonging to each group are shown above each violin. In panel A, the dotted line represents a threshold of 0.85 and in panel B, the dotted line represents a threshold of 0.9; details of thresholds discussed in main text). **C and D**. Column plot showing the proportion of references (y-axis) from each human labeled category (“reporting an occurrence”, “maybe reporting an occurrence”, or “not reporting an occurrence”; x-axis) that would be retained if a threshold small language model score was used for filtering references. The proportion of each column in the positive y space indicates the fraction of references that would pass the filter and be retained, while the proportion of each column in the negative y space indicates the fraction of references that would be rejected by the filter and eliminated. Exact proportions are shown in numbers above and below each column. In panel C the threshold is 0.85, based on two-prompt scoring, while in panel D the threshold is 0.9, based on single, general prompt scoring (details in main text and methods section). For example, if a score of 0.85 were used as a threshold with which to filter references that had been scored using the two-prompt small language model scoring system, then 86% of references reporting occurrences would be retained while 14% of such references would be rejected, 35% of references maybe reporting occurrences would be retained, while 65% of such references would be rejected, and 20% of references not reporting occurrences would be retained while 80% of such references would be rejected. In all panels A-D, colors correspond to the three human label categories (“reporting an occurrence”, “maybe reporting an occurrence”, “not reporting an occurrence”). BART stands for the bart-large-mnli small language model.

Next, we investigated the ability of these scores to act as a filter to separate articles of interest that report chemical occurrences from those that did not report such occurrences. Thus, we examined the proportion of the former type articles that would be retained if a threshold score were to be used as a filtering criterion for the reference collection (i.e., if references with a score higher than a threshold were to be retained and those with a score lower than the threshold were to be eliminated from the collection).Based on the distribution of scores assigned to articles that reported chemical occurrences versus those that did not (**Fig. 2A and B**), we selected 0.85 as a threshold for the specific two-prompt scores and 0.90 as a threshold for the general prompt-derived scores. With these thresholds, the specific two-prompt scoring system acting as a filter would have retained 86% of the references that report phytochemical occurrences (the references of interest in our study), and rejected 80% of the references that did not report an occurrence (**Fig. 2C**). The general prompting system, with a 0.90 filtering threshold, would have retained 92% of the references reporting phytochemical occurrences and eliminated 55% of the references that did not report occurrences (**Fig. 2D**). While both the two-prompt and general prompt filtering approach led to the retention and rejection, respectively, of article of interest and not of interest, the two approaches handled articles we had labelled as “maybe reports an occurrence” differently: the two-prompt approach kept only 35% of these, while the general approach kept 81%. To learn more about these “maybe” references, we obtained and read 100 full text articles for these references (those related to α-amyrone and dammarenediol II, **Supplemental File 3**). This manual inspection revealed that approximately 65% percent of these “maybe” references contained reports of compound occurrence data, which suggested that access to full text information will help create more comprehensive chemical occurrence datasets. However, regardless of whether full texts are available or not, our results show that small language model relevance scores provide a means to quickly (∼45,000 references / hr.) and accurately (∼80% relevant articles kept, ∼80% of irrelevant articles rejected) identify references that are most likely to provide the information that a user might be seeking. This ability will be highly useful when dealing with many thousands of references. Our data also indicate that there will likely be a benefit to developing more nuanced filtering approaches to handle edge cases like the ‘maybe’ articles we identified here.

### 2.2: SLM Task B: Extracting compound occurrence data with language models

After filtering our collection of references to include only entries with high scores concerning phytochemical occurrence data, we evaluated the ability of language models to extract experimentally supported compound presence details. In this task, two steps can be envisioned: (i) a first step in which a model receives a body of text including the title and abstract of a scientific article and (ii) a second step in which a model receives a query about compound occurrences. For example, in the second step, we might ask the model: “Does the provided text offer experimental evidence that *Arabidopsis thaliana* produces the chemical compound thalaniol?” This mode of operation represents a **targeted approach**. A second mode of operation (for the second step) could be to pass a language model a text passage containing the title and abstract of a scientific article and pose an open-ended query such as: “List all of the plant species mentioned in the provided text and indicate which chemical compounds were reported from each one as part of the experimental investigation described in the passage.” This second mode represents an **untargeted approach**. Several advantages and disadvantages of each approach can be imagined from the outset. For example, an untargeted approach does not require a preconceived set of chemical compounds or plant species of interest about which to query the model, and a single untargeted query can potentially extract multiple compound-occurrence data simultaneously. In contrast, one benefit of the targeted approach is the relative simplicity of creating human-labeled data. Thus, a true/false answer about one plant/compound occurrence can be supplied by the human or model instead of meticulously generating a complete list of such occurrences. Furthermore, a model’s rate of detecting true negative associations can be measured directly by comparing the model’s response to plant and compound names appearing in an abstract, without experimental association data, to the corresponding human response. To summarize, the targeted and untargeted approaches each offer distinct benefits. Therefore, we tested and herein present results from both approaches.

#### 2.2.1 SLM Task B1: Targeted compound occurrence data extraction

To evaluate the ability of large language models to extract compound occurrence data from scientific abstracts, we first prepared and manually evaluated a set of candidate occurrences. For this effort, we used regular expression-based pattern matching to identify accepted plant species names in the abstracts associated with the six triterpenoids that comprised the present test case. We then compiled a data set containing three columns: the title and abstract of each reference, the chemical compound linked to it (the SciFinder® search compound that retrieved that reference in the first place), and accepted plant species name(s) found in that title or abstract. We manually evaluated 500 candidate associations and annotated each occurrence as positive (the abstract described experimental support for the occurrence of that compound in that plant species) or a negative (the abstract did not provide such support). We found that roughly 350 (71%) of the candidate associations were negatives, while around 150 (29%) were positives (**Supplemental File 4**). With a set of human-labeled compound species or candidate compound species associations in hand, we next turned to evaluating whether open-source language models could perform the same task. For this task, we used open-source language models that accepted two types of prompts. The first prompt was a system prompt that contained detailed instructions on how the model should generate an output. The second prompt (also called user text) delivered content from which the model generated that output. We used the second prompt to supply information on the candidate compound species association (title/abstract, compound name, and species name) and the system prompt to convey detailed instructions on how the model was supposed to evaluate this given information (full details in Methods).

Past research has shown that language models of different sizes vary in their ability to perform natural language processing tasks(Brown et al. 2020; Kaplan et al. 2020), including tasks related to chemical occurrence data extraction(Busta et al. 2024a). Accordingly, in evaluating their capacity for the present targeted occurrence extraction task, we tested 12 language models of various scales, spanning 0.5 billion to 32 billion parameters (often denoted 0.5B to 32B, **Fig. 3A**). These models included variants of different sizes from the Qwen family (Qwen: An et al. 2025) (32B, 14B, 7B, and 0.5B), the Gemma family(Gemma et al. 2025) (27B, 12B, 4B, and 1B), and the Phi- 4 family (phi-4 14B and phi4-mini-instruct 4B)(Abdin et al. 2024). Each model was given the same system prompt and all 500 candidate occurrences that had been previously examined manually. During these assessments, all models were run at 16-bit precision, except gemma-3-27B-it-unsloth and phi-4-unsloth-bnb-4bit, which are dynamically quantized instances operating at 4-bit precision (**Fig. 3A**). When reviewing the 500 candidate associations, run times generally varied inversely with size; qwen-2.5-32B-instruct handled about 200 references per hour, while qwen-2.5-0.5B-instruct surpassed 32,000 per hour (**Fig. 3A**). Notably, the quantized variants processed references at speeds only slightly higher than their full-resolution counterparts (for example, the 4-bit phi-4-unsloth at 1,500 references / hr. and the 16-bit phi-4 at 1,200 per hour). These speeds will be important when applying language model-based approaches to larger projects or the assembly of databases.

**Figure 3:**
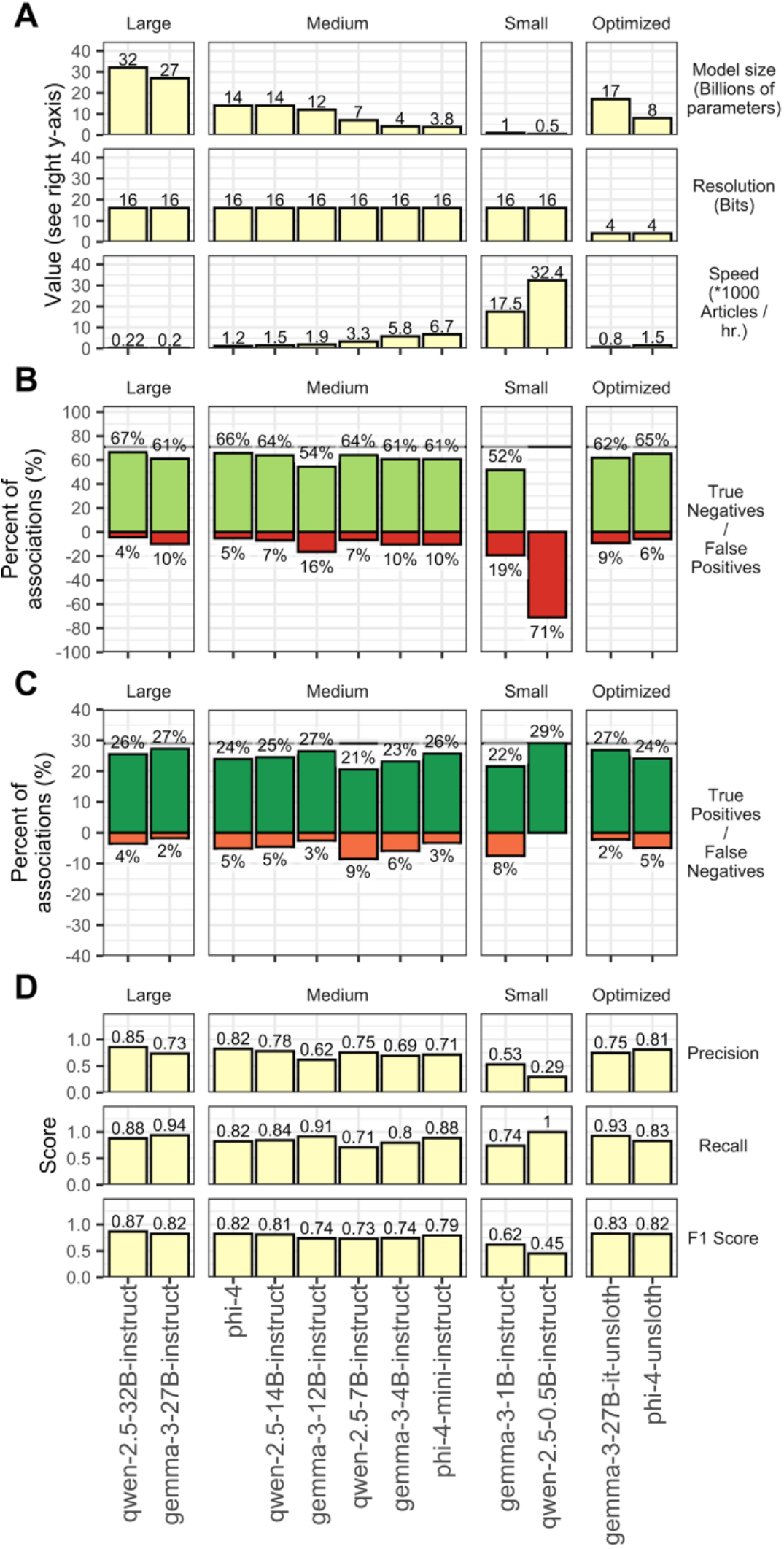
Performance of language models on a targeted compound occurrence data extraction task. **A**. Bar plot showing various metrics (y axes in each row of panels) for different language models (x axis). The first row shows model size in billions of parameters, the second row shows model resolution in bits, the third row shows the speed with which a model processes references (using the prompt shown in the methods section) in units of 1000 references per hour. **B**. Bar plot showing the raw performance metrics of each model (false negative, false positive, true negative, and true positive rates). False negatives arise when a model erroneously marks a real compound occurrence as not being true. False positives arise when a model erroneously marks a simple textual occurrence of a compound name and species name as an occurrence data point. True negatives arise when a model correctly marks a simple textual occurrence of a compound name and species name as such, and not as an occurrence data point. True positives arise when a model correctly marks a compound occurrence as such. According to human evaluation of the 500 putative occurrences used to test the models, 71% of the putative occurrences were real (i.e. “positives”), and 29% of the putative occurrences were just textual co-occurrence (i.e. “negatives”). Thus, a perfect model would have, in this experiment, a 71% true negative rate and a 29% true positive rate. Bars are colored according to true/false positive/negative. **C**. Bar plot showing the processed performance metrics of each model. In the first row, the precision of each model is shown (the ratio of true positives to the sum of true positives and false positives). In the second row, the recall of each model is shown (the ratio of true positives to the sum of true positives and false negatives). In the third row, the F1 score is shown, which is the harmonic mean of the precision and recall. In A–C, models are organized into columns of panels by type (large: > 20 B parameters, medium: 1–20 B parameters, small: 0–1 B parameters, and optimized: 4-bit resolution models).

Alongside measuring how quickly various models processed 500 candidate associations, we also examined model accuracy. To gauge that accuracy, we compared whether each model labeled every candidate association as positive or negative against the corresponding human label. The results let us classify each model output as a true positive (when the model labeled a candidate association as positive, matching the human label), a true negative (when both the model and the human labeled it negative), a false positive (when the model labeled it positive but the human did not), or a false negative (when the model labeled it negative but the human did not). Because 71% of the 500 candidate associations were negative, a high-performing model would have a true negative rate approaching 71%. The true negative rates for the models tested ranged from 52% to 67%, with models containing more parameters generally showing higher percentages (**Fig. 3B**). One exception was qwen-2.5-0.5B-instruct, which had a 0% true negative rate, as it labeled all candidates occurrences as positive. These differences in true negative rates came with parallel differences in false positive rates, since false positives arise when a model incorrectly labels a negative result as positive. The false positive rate is one of the most important metrics for this task because those errors represent fabricated occurrence data. In our experiments, larger models achieved lower false positive rates overall, with qwen-2.5-32B-instruct and phi-4 showing the lowest values at 4% and 5%, respectively (**Fig. 3B**). Because both were also the slowest and largest, there is a clear trade-off between parameter count and computational requirements on one hand and task-specific accuracy on the other.

The models we tested here did not only vary in their (true negative)/(false positive) rates, but also in their (true positive)/(false negative) rates. Since positive associations comprised 29% of the 500 candidate associations, a perfect model in our experiment would have a 29% true positive rate. True positive rates among the models tested here generally ranged from 21% to 27% (**Fig. 3B**). This variability did not correlate as strongly with model size as did the (true negative)/(false positive) rates. For example, a large model (qwen-2.5-32B-intruct), two medium models (gemma-3-12B-instruct and phi-4-mini-instruct (4B)), and one of the quantized models (gemma-3-27B-it- unsloth) all had very similar true positive rates (26% or 27%, **Fig. 3B**). Note that the perfect true positive rate of qwen-2.5-0.5B-instruct is a misleading statistic, since this model simply labeled all associations with which it was presented as positive. To account for such potentially misleading rates, we computed precision and recall statistics. Precision is calculated as the number of true positive results divided by the sum of true positive and false positive results, which indicates how reliable the model is when it marks an association as positive. Recall is calculated as the number of true positive results divided by the sum of true positive and false negative results, which reflects the model’s ability to correctly identify all actual positive associations. Excluding qwen-2.5-0.5B-instruct, precision varied from 0.5 to as high as 0.85 and recall varied from 0.71 to 0.94 (**Fig. 3C**). We also computed F1 scores, which are the harmonic mean of precision and recall, to provide a single metric to balance both reliability (precision) and completeness (recall). F1 scores (excluding qwen-2.5-0.5B-instruct) ranged from 0.62 (gemma-3-1B-instruct) to 0.87 (qwen-2.5-32B-instruct) and varied, again, according to model size, which reinforced the importance of that parameter in task-specific accuracy.

So far, our results indicated that language models can assess whether an abstract describes experimental support for a particular compound, but no model was entirely accurate in performing this task. Accordingly, we next turned our attention to a detailed examination of the candidate associations that were frequently labeled incorrectly by the language models. Specifically, we reviewed the incorrect answers generated by the phi-4 model. First, we focused on references in which no experimental support for a compound’s occurrence was provided, yet the model (erroneously) indicated such support was presented (i.e., false positives). Among these occurrences, two main text structures appeared to “confuse” the model. The first scenario involved abstracts where occurrence data were not presented in separate sentences but instead merged with multiple data types. For example, some passages combined information from authentic standards and plant extracts, or from sediments and plant extracts, or listed multiple compounds from several species in a single statement. The second scenario leading to false positives involved abstracts that failed to provide clear statements about plant/compound occurrences, even to a human reader. As an example, one such abstract stated “beta-sitosterol and alpha-amyrin were isolated from unsaponifiable fractions of mature seeds of solanaceae plants” and mentioned the solanaceous species *Hyoscyamus muticus*, which caused the model to label alpha-amyrin as present in *Hyoscyamus muticus*, even though this link was not explicitly supported by the text. Finally, we examined references where positive associations were mistakenly labeled by the models as negative (i.e., false negatives). We identified three main cases: (i) abstracts that were written in confusing ways, which lead the model to produce an incorrect result, (ii) abstracts that contained an alternative spelling or abbreviation for a compound or species name, and (iii) clearly written abstracts in which the model nevertheless failed to provide the correct answer. These scenarios appeared in roughly equal proportions among phi-4’s false negatives. To summarize, the model sometimes makes clear mistakes, but, just as often, the model produces incorrect answers because of inconsistencies or unclear information in the input data. Finally, we also examined the performance of the models when alternative spellings of compound names were present in abstracts. Across the 500 candidate associations we manually evaluated there were 28 instances where alternative spellings were used in the abstract (amyrin/amirine, friedelin/friedeline, amyrone/amyrenone). Evaluating these candidate associations, the highest performing models were correct ∼50% of the time, which is lower than model performance across the entire dataset (∼10% overall error rate). Thus, we conclude that these alternative spellings do impact model performance and strategies to deal with such should be included in the design of small language model-based pipelines.

Several conclusions arise from our work with targeted compound-occurrence data set extraction. First, models with more parameters (“larger” models) appear to perform the task with higher accuracy, though that improvement comes alongside increased computational demands and time requirements. To balance speed and performance, architectures such as phi-4 stand out from those evaluated in this study. Next, the abilities of systems like phi-4 to accurately detect true negatives indicate that they are distinguishing references with textual co-occurrence of plant and compound names from references that present experimental evidence for a plant producing a given compound. Finally, our examination of the underlying reasons for incorrect answers revealed many errors arise from inconsistencies or unclear information in the input data, which suggests that using full-text articles instead of titles and abstracts may improve results beyond the approach described here.

### 2.2.2 SLM Task B2: Untargeted compound occurrence data extraction

After assessing the extent to which language models can classify compound occurrences in a targeted manner, we next examined these systems’ abilities with the same task in an untargeted way. For this process we used models that, as before, accept a system prompt with detailed instructions and a second prompt containing content with which to work. Our general approach was to provide a system prompt directing the model to read the input text (title/abstract) and write all experimentally supported compound occurrences in a Python dictionary format (for example: {“Arabidopsis thaliana”: [“arabidiol”, “beta-sitosterol”], “Brassica oleracea”: [“beta-sitosterol”, “alpha-amyrin”]}). Thus, this task is considerably more complicated than the targeted approach. Due to this complexity, we conducted some preliminary tests to determine which of our 12 models might be suitable for this task. We found that the two large models and the two small ones were, respectively, too slow and too inaccurate to be feasible. For this reason, we proceeded with the six medium models as well as the two quantized 4-bit variants described in the previous section. Previous work has shown that the exact phrasing of system prompts can have substantial impacts on the accuracy of language model outputs (Razavi et al. 2025; Sclar et al. 2024), which included the context of phytochemical data processing (Knapp et al. 2024b). This phenomenon is the basis for prompt engineering. This untargeted task was inherently more complicated than the above-described targeted approach, but further complications arose because we wanted a specific output format (the Python dictionary). We investigated a variety of prompts to determine how they might impact results from each model. As in the previous sections, to benchmark the ability of the models to perform this task, we again began by performing this task manually. We read 100 abstracts and wrote out the compound species associations reported in each in the JSON, or Python dictionary, format. This led to the identification of just over 400 compound occurrences across the 100 abstracts (**Supplemental File 5**). Below, we describe the performance of the 8 models and the 11 prompts on this untargeted compound occurrence extraction task with the 100 manually evaluated abstracts.

To begin, we carefully created a detailed system prompt and then employed a commercial large language model to produce 10 additional prompt variants that contained the same instructions but with different phrasings (all prompts included in **Supplemental File 6**). We then used each of the eleven prompts to instruct each of the eight models to write out all experimentally supported occurrences in each of the 100 manually evaluated abstracts. Next, we examined the ability of each model/prompt combination to provide results in a valid Python dictionary (the structure of the response needed to perform this data extraction task) and the speed at which each model/prompt combination could process the 100 abstracts. The percentage of responses from each model in answer to each prompt varied considerably, with some model/prompt combinations producing zero valid dictionaries and others generating 100% valid dictionaries (**Fig. 4A**). Most model-prompt combinations produced >90% correctly structured responses, with some notable exceptions. Interestingly, qwen-2.5-14B-instruct struggled to consistently produce valid dictionary outputs, while its smaller sibling, qwen-2.5-7B-instruct, yielded over 90% valid dictionaries in most cases. This result breaks the trend of larger models being more proficient, as described in the previous section of this report. Phi-4 was the best model tested at this task since it returned 100% valid Python dictionaries, except for one response to prompt 8 (**Fig. 4A**). We also observed variation among the prompts tested, with prompts 9, 10, and 4 eliciting higher proportions of valid responses across all the models than other prompts. We also examined the rate at which each model and prompt pairing could process queries. Rates ranged from about 200 references per hour to almost 1,500 references per hour, with model size as the primary determinant of speed (**Fig. 4B**). Different prompts sometimes caused variability in processing times for the same model, though these shifts were negligible compared to those driven by scale. Overall, the largest model, gemma-3-27B-instruct-unsloth, was the slowest. Meanwhile, phi-4-mini-instruct and qwen-2.5-7B-instruct performed the fastest, at rates around 1,000 articles or references per hour. Altogether, the outcomes suggested that the phi-4 family models, along with qwen-2.5-7B-instruct combined with prompts 9, 10, and 4, were the most accurate for further detailed investigation. The four best performing models for producing valid Python dictionaries included the two fastest frameworks (phi-4-mini-instruct and qwen- 2.5-7B-instruct), which showed that larger models do not always perform more proficiently than smaller versions.

**Figure 4:**
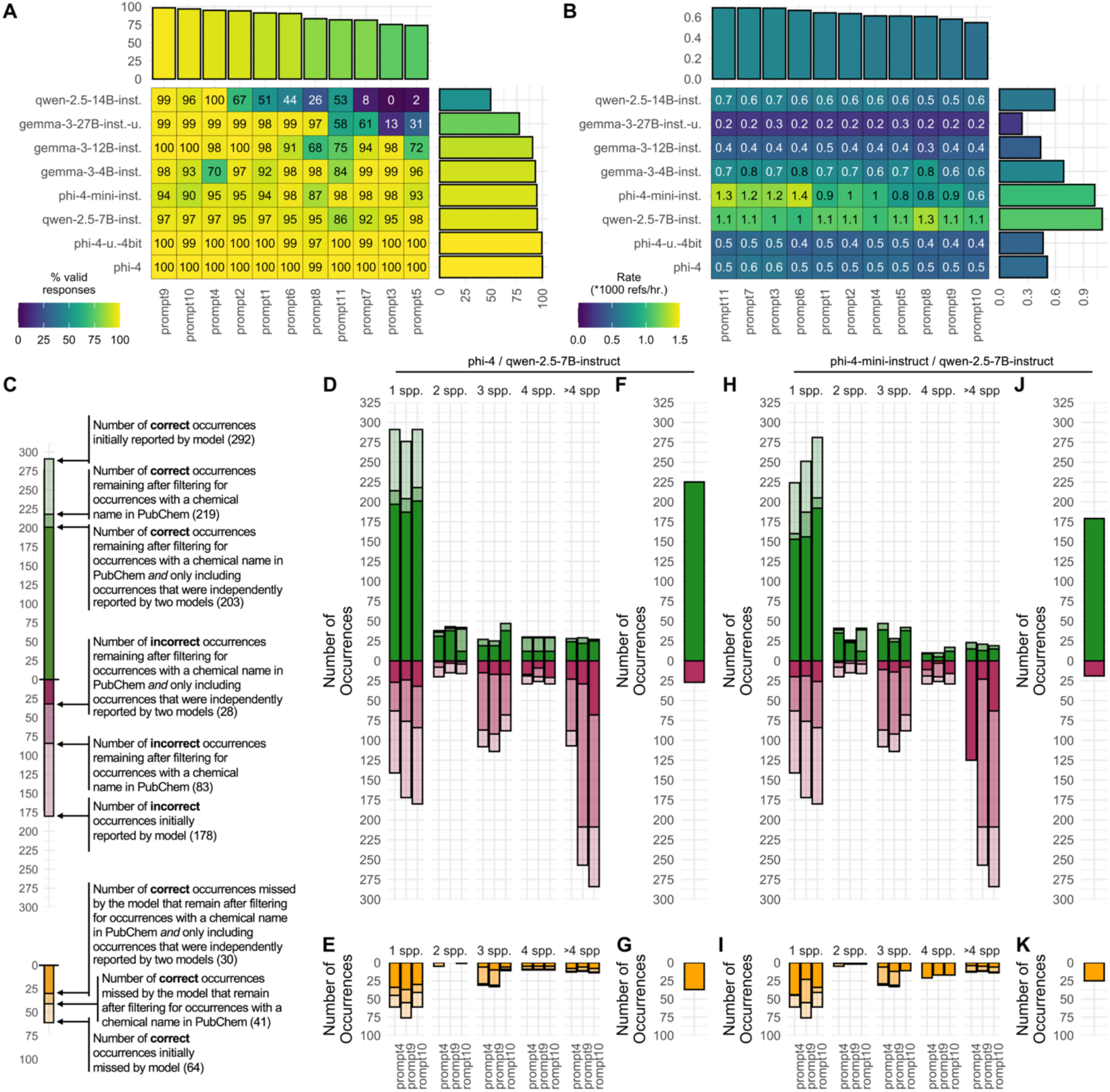
Performance of language models on a targeted compound occurrence data extraction task. **A**. Heat map showing the percent of outputs that contain valid python dictionaries (encoded with color and written inside each box) from each language model (y-axis) in response to each prompt (x-axis). The marginal (i.e. top and right) plots show the mean percent valid responses across all models for each prompt or across all prompts for each model. **B**. Heat map showing the rate (in 1000 references per hour) of processing by each language model (y-axis) in response to each prompt (x-axis). The marginal (i.e. top and right) plots show the mean percent valid responses across all models for each prompt or across all prompts for each model. **C**. Guide describing how to interpret panels D-K. **D-K**. Evaluation of occurrence data reported by language models (D/E/F/G: phi-4 and, in darkest bars, phi-4 in agreement with qwen-2.5-7B-instruct; H/I/J/K: phi-4-mini-instruct, and, in darkest bars, phi-4-mini-instruct in agreement with qwen-2.5-7B-instruct. D and H show the number of correct occurrences (true positives, positive y-axis) and incorrect occurrences (false positives, negative y-axis) reported, as indicated in panel C. E and I show the number of correct occurrences (false negatives, negative y-axis) reported, as indicated in panel C. F and J show the number of correct occurrences (true positives, positive y-axis) and incorrect occurrences (false positives, negative y-axis) reported after filtering for occurrences whose compounds are in PubChem and were agreed upon by the two models. G and K show the number of correct occurrences missed by the models after PubChem and agreement filtering (false negatives, negative y-axis).

In the previous section, we identified that the results from prompts 9, 10, and 4, in conjunction with phi-4, phi-4- mini-instruct, and qwen-2.5-7B-instruct warranted further scrutiny. Therefore, we next examined the accuracy of occurrences generated by those models in response to those prompts. In contrast to our quantitative assessment of the models’ ability to evaluate targeted compound instances, this broader approach allowed for quantifying only three response types: true positives (correct occurrences reported by a model), false positives (incorrect occurrences reported), and false negatives (correct occurrences missed by the model but found during manual evaluation, **Fig. 4C**). Note that true negatives are not present in this untargeted analysis since the model is only asked to report existing occurrences, not to classify candidate occurrences. We quantified the number and category of each occurrence identified by each model in response to prompts 4, 9, and 10. We observed that using different system prompts led to only minor variations in the total correct versus incorrect instances flagged by a given model, but, interesting, that correct versus incorrect outputs varied greatly with respect to the number of species described in a given abstract (**Fig. 4D, 4E, 4H, and 4I**). Specifically, references involving more than four species appeared “confusing” to the models, resulting in large numbers of inaccuracies from those sources (**Fig. 4D and 4H**), while abstracts focused on one or two species typically yielded substantially more correct instances compared to incorrect ones (**Fig. 4D and 4H**). Even so, the ratio of correct to incorrect responses typically generated from articles reporting on one or two species was roughly 2:1, an approximately 30% false positive rate.

To reduce the false positive rate observed during this untargeted compound occurrence extraction task, we introduced two types of filters. For the first filter, we programmatically compared the compound name reported in each occurrence against the PubChem database to check if it appeared among the entries. We removed all reported occurrences describing compounds missing from PubChem, which generally produced a bigger drop in incorrect results than in correct ones. The second filter relied on two language models identifying the same occurrence from a given abstract. Only those occurrences found by both, working independently, were kept, while partial matches (instances flagged by a single model but not recognized by another) were excluded. We tested this two-part filtering approach with two pairs of models: (i) one containing the most advanced model: phi-4 + qwen-2.5-7B-instruct, and another featuring the two fastest options: phi-4-mini-instruct + qwen-2.5-7B-instruct. In both scenarios, the agreement filter yielded a marked decrease in inaccurate entries in the final dataset and only a small decline in valid ones (**Fig. 4D and H**). Finally, to produce a dataset that reflects the lowest likely false positive rate for these models on the untargeted task at hand, we combined three filtering strategies: we restricted data to abstracts mentioning one or two species, retained only occurrences describing chemicals found in the PubChem database, and kept only those occurrences that were independently detected from the same abstract by two different language models. Using this threefold approach, phi-4 + qwen-2.5-7B-instruct produced about 225 accurate occurrences and 25 inaccurate ones (an 11% error rate and ∼55% yield, relative to the 400 occurrences found during manual inspection of the 100 abstracts, **Fig. 4F**). Meanwhile, phi-4-mini-instruct + qwen-2.5-7B-instruct yielded 175 valid occurrences and around 20 erroneous findings (also an 11% error rate and ∼44% yield/recall, **Fig. 4J**). Thus, pairing two fastest models led to a dataset that was less comprehensive but maintained similar accuracy as a pair that contained a considerably larger and more sophisticated model.

## 3. Conclusions and Future Directions

Here, we evaluated the ability of small language models to perform two major tasks: to numerically score references based on their relevance to a given topic (SLM Task A) and to extract structured data from unstructured inputs in both a targeted (SLM Task B1) and untargeted fashion (SLM Task B2). Our efforts showed that a small language model could rapidly and effectively score references in a set so that a threshold score could be used as a filter to substantially enrich that set for articles of interest (Section 2.1). Limitations arose when handling edge cases, with highly tailored, task-specific prompts emerging as a possible approach to address those shortcomings. When using small language models to classify candidate compound occurrences as true or false, we observed that if an abstract directly reports the detection of a particular compound in a specific plant species, the models nearly always label the candidate occurrence correctly (Section 2.2.1). In this task however, a trade-off did appear between accuracy and model size (parameter count) and compute requirements. Among the mistakes noted (false positives as low as 5% and false negatives as low as 2% for certain models), these misclassifications were as often tied to convoluted and unclear writing in the input abstract as they were to outright model errors. For extracting compound occurrence information from unstructured text in an untargeted manner, we found small language models to be effective, though choosing a suitable model and pipeline strategy proved more challenging than earlier tasks (Section 2.2.2). We discovered that prompt engineering, selecting a model, and filtering reported detections by cross-referencing chemical databases, along with requiring two small language models to independently agree on an occurrence, yielded the best reporting statistics (∼10% false positives and ∼50% yield). Of note, this relatively low yield arises because many correct associations are filtered out, essentially sacrificed, to lower the false positive rate. Regarding all tasks considered, more advanced prompting techniques (e.g., chain-of-thought prompting (Wei et al. 2022) or model distillation (Hinton et al. 2015; Sanh et al. 2020) could reduce error rates further and improve yield/recall. In addition, future model releases, including small reasoning models, may also address these limitations. Finally, we will note that many abstracts we worked with here presented problems for humans and language models alike by failing to contain clear and concise information. We read hundreds of abstracts for the present project. Fully understanding many abstracts in a timely fashion was extremely difficult due to long, convoluted sentences, the presentation of connected data types (e.g., plants and compounds) in multiple sentences spread throughout a long abstract, the use of compound numbers or abbreviations instead of compound names, poor grammar, and so forth. In a variety of cases, we were surprised that the language models performed reasonably well while humans needed considerable time to understand the same abstracts.

Overall, though the approaches here represent a considerable advance over manual curation (at least, with respect to the creation of large databases, where speed is a prime consideration), a substantial amount of plant chemical occurrence data will still not be retrieved from the literature using the techniques presented here. One important step forward will be the development of pipelines that can handle articles reporting occurrence data from dozens of species, including in tabular format. In addition, further attempts towards occurrence databases, and in fact scientific endeavors in general, need literature databases that include the full text files along with reference citations and abstracts. The separation of the full text from the citations seems to be a systematic and legal barrier that needs to be overcome. The expanded posting of pre-prints is suggested as a potential, albeit partial, solution to this issue. In addition to the tasks we quantitatively evaluated here, we also experimented with several versions of Microsoft’s Phi-4 model to conduct multiple activities related to reference citations (e.g., species name extraction, compound number or plant number extraction, etc.) and found that the models could perform a range of additional functions, suggesting versatility and application in order domains. In our case, these functions have allowed us to identify publications that most likely contain extensive tabular data in the full text, flagging them for analysis by a pipeline suitable for such reports. Finally, in our efforts, we found that filtering capabilities such as those provided by SciFinder® and EndNote^TM^ showed usefulness in a somewhat orthogonal way to the value of the small language model scores. For example, in our case, we were able to eliminate many articles of low relevance to our case studies using EndNote™ keyword filters. As these commercial software tools and other related programs are outfitted with language model (“artificial intelligence”) capabilities, it will be important to evaluate and incorporate those features into discipline-specific workflows. We strongly encourage the scientific community to look for new versions of their favorite research tools that incorporate language model features, and to experiment and empirically test and report on such functionality in field-specific tasks as they emerge.

## 4. Methods

Literature searches were conducted with CAS SciFinder®. SciFinder® searches were conducted by entering the compound CAS Registry® number from the SUBSTANCE menu and then working with all the references that were assigned to this Registry number. SciFinder® references were downloaded as “tagged” text files. The “tagged” text file selection provides numerous fields including the CAS Registry numbers for all compounds discussed in a given article. Multiple tagged files were downloaded for each compound (according to year ranges) since the SciFinder® software limits an individual tagged export file to 100 citations. SciFinder limits the number of citations that can be exported in one file to 100. Thus, for a compound such as alpha-amyrin with 4,344 SciFinder references, the downloading of all references was not possible. If the number of filtered references was greater than 400, the word “plant” was entered into the “search within results.” Thus, only English-language journal references that corresponded to the “search within results” term “plant” were downloaded (1,744 references, in the example of alpha-amyrin). PubMed® searches for the six triterpenoids were also conducted based on their major common names (not all synonyms were used). These PubMed® searches were conducted with the compound names shown at the top of Table 1 since PubMed® does not generally recognize CAS Registry® numbers. PubMed® files were downloaded as PubMed (NLM) files. Of note is that PubMed® provides automated access to its search and abstract download services through a REST API and various language-specific packages like R and Trez. These tools could be leveraged in the future to further streamline literature analysis projects and automate data extraction and tabulation.

EndNoteTM Version 21.5 (https://endnote.com/) was used to import and combine the sets of “tagged” SciFinder® export text files for each compound into an individual EndNoteTM compound folders (with the “discard duplicate” feature turned on). Furthermore, EndNoteTM “Smart Groups” were set up for each of the six triterpenoids, which included the CAS Registry® number and multiple names for each compound (i.e., synonyms). The references in each of the six Smart Groups were then added to the corresponding original six triterpenoid folders (with automatic elimination of duplicates). As noted above, some plants contained more than one of the six triterpenoids. These EndNoteTM operations ensured that any references that might have been missed in a given SciFinder® compound search, but included in another compound search, would end up in the appropriate folders (i.e., one reference might be in more than one compound folder). In EndNote®, the user can select scores of references and then right-click on “Find full text.” EndNote will then automatically download the PDF files for each reference that cites a journal for which the user’s institution has a subscription or an open-source journal. However, in some cases, software blocks (e.g., the “Are you a human filter?”) prevent the downloading of some files. In our case at our institution, EndNote downloads approximately 40-50% of the PDF files for the selected references.

All manual evaluation of reference relevance (“reports an occurrence”, “maybe reports an occurrence”, “does not report an occurrence”), manual evolution of candidate occurrences (targeted) and manual extraction of associations (untargeted) was performed by opening the list of references in Microsoft Excel and entering the manual annotations into a new column. References were labeled as “maybe reports an occurrence” if they mentioned specific plant species and the isolation of multiple compounds from the species but did not mention the specific compounds’ names in the abstract. While the likelihood of a plant/compound association appearing in the full article was high, we nevertheless conservatively chose to label these types of citations as “maybe reports an occurrence” until the full text article file could be evaluated. An “maybe” example is: “Medicinal attributes of Solanum capsicoides All.: an antioxidant perspective. Int. J. Pharm. Sci. Res. 12(5): 2810-2817. The study evaluates the medicinal efficacy of Solanum capsicoides fruits as an antioxidant. Fruit extracts were prepared using acetone, ethanol, HCl, and water […] A neg. correlation was observed between the pigments, anthocyanins, and carotenoids, with DPPH and CUPRAC activity. […] From this study, it can be considered that the phenolics present in the fruits contribute to the characteristic antioxidant property.”

The Facebook BART-Large-MNLI zero-shot classification model (https://huggingface.co/facebook/bart-large-mnli) was applied to the individual sets of compound reference citations in the EndNoteTM database. The model was run on a single NVIDIA GV100GL [Quadro GV100] GPU. First, the set of references in the curated EndNoteTM folder for a given compound was selected and exported from this folder to a text file (with the “annotated” style selected). This text file was then imported into an Excel file (e.g., with the legacy “get text from file” Excel wizard. The resulting Excel sheet was then modified so that each reference citation (author/year/journal/abstract) was contained in one cell and all cells resided in one column. This Excel sheet, which contained all the reference citations for a given compound, was then saved as a CSV UTF-8 (Comma delimited) file. This CSV file was used via JupyterLab (https://jupyter.org/, operating in a WINDOWS 11 environment) and a custom Python program (full code in Supplemental File 8). System prompt-accepting chat language models were downloaded from HuggingFace.co and run on a single NVIDIA GV100GL [Quadro GV100] GPU using custom code (full code provided in Supplemental File 8). Calculation of precision, recall, and F1 scores as well as plotting were performed in R. Additional system prompts for the prompt engineering reported in Section 2.2.2 were generated by OpenAI’s 04-mini-high language model using the ChatGPT browser interface.

## 5. Supplemental Materials

Supplemental File 1: 1,558 references manually scored for relevance to compound occurrence.

Supplemental File 2: Details of classifier phrases.

Supplemental File 3: Details of maybe references.

Supplemental File 4: 500 manually evaluated candidate occurrences.

Supplemental File 5: 100 abstracts from which untargeted occurrence data was manually extracted.

Supplemental File 6: Prompts that were used in small language model untargeted occurrence data extraction.

Supplemental File 7: Schematic of reference acquisition process.

Supplemental File 8: Code used in this work.

## 6. Acknowledgements

The authors wish to acknowledge the support the University of Minnesota Duluth Chemistry and Biochemistry Department. During the writing of this manuscript, the large language model o4-mini-high from OpenAI was utilized for copy editing to suggest alternative sentence structures and word choices that, in our view, enhanced grammar and readability. Finally, we collectively acknowledge that the University of Minnesota Duluth is located on the traditional, ancestral, and contemporary lands of Indigenous people. The University resides on land that was cared for and called home by the Ojibwe people, before them the Dakota and Northern Cheyenne people, and other Native peoples from time immemorial. Ceded by the Ojibwe in an 1854 treaty, this land holds great historical, spiritual, and personal significance for its original stewards, the Native nations, and peoples of this region. We recognize and continually support and advocate for the sovereignty of the Native nations in this territory and beyond. By offering this land acknowledgment, we affirm tribal sovereignty and will work to hold the University of Minnesota Duluth accountable to American Indian peoples and nations.

## 7. Data Availability Statement

All data and code used in this study are available for free as a Supplement to this document.

