## Supplemental File 2 for "Small language models enable rapid and accurate extraction of structured data from unstructured text: an example with plants and their specialized metabolites"

### Alpha-Amyrin (CAS 638-96-9)

#### Compound names

Multiple names for alpha-amyrin appear in the CAS Registry® (SciFinder®). These names are listed below along with the number of occurrences in the EndNote database (out of 231 total alpha-amyrin citations):

- α-amyrin
- (3β)-Urs-12-en-3-ol
- Urs-12-en-3β-ol
- Viminalol
- 3β-Hydroxyurs-12-ene
- α-Amyrenol
- α-Amyrine

The names along with the number of occurrences in the EndNote database (out of 1,341 total alpha-amyrin citations):

- Amyrin: 630 titles or abstracts
- Urs-12-en-3-ol: 2 titles or abstracts
- Viminalol: 2 titles or abstracts
- Hydroxyurs-12-ene: 3 titles or abstracts
- Amyrenol: 1 title or abstract
- Amyrine: 6 titles or abstracts

The total occurrences of these compound names were 645 out of a total of 1,341 references. The remaining references may still be **OF-INTEREST** but lack the target name in the abstract or title.

#### SLM classifier phrases

The names, other than "amyrin" only amounted to 14 references (1%) of the total references. Since the SLM seemed to work better with fewer names, the following phrases were used for the small language model run:

- "Amyrin is present in plants."
- "Amyrin is not present in plants."

#### Summary statistics or alpha-amyrin

- Total number of references in EndNote database = 1,341.
- Total number of references that contained CAS Registry names in title and/or abstract = 645

The 118 citations with the highest SLM scores (0.992 – 0.998) were ranked by a human. The results were:

- Total **OF-INTEREST** references* = 97
- Total **POTENTIALLY-OF-INTEREST** references* = 10
- Total **NOT-OF-INTEREST** references* = 11
  - *According to human evaluation of the titles and abstracts for each citation.

### Alpha-Amyrone (CAS 638-96-0)

#### Compound names

Multiple names for alpha-amyrone appear in the CAS Registry® (SciFinder®). These names are listed below along with the number of occurrences in the EndNote database (out of 231 total alpha-amyrone citations):

- α-amyrone: 16 titles or abstracts
- urs-12-en-3-one: 9 titles or abstracts
- 3-ketours-12-ene: 0 titles or abstracts
- 3-oxours-12-ene: 0 titles or abstracts
- α-amirenone: 1 abstract
- α-amirone: 0 titles or abstracts
- α-amyrenone: 44 titles or abstracts

The total occurrences of these compound names was 70 out of a total of 231 references. The remaining references may still be **OF-INTEREST** but lack the target name in the abstract or title.

#### SLM classifier phrases

The following phrases were used for the small language model run:

- "Amyrone, urs-12-en-3-one, amirenone, or amyrenone is present in plants."
- "Amyrone, urs-12-en-3-one, amirenone, or amyrenone is not present in plants."

#### Summary statistics or alpha-amyrone

- Total number of references in EndNote database = 231.
- Total number of references that contained CAS Registry names in title and/or abstract = 70
- Total **OF-INTEREST** references* = 53
- Total **POTENTIALLY-OF-INTEREST** references* = 73
- Total **NOT-OF-INTEREST** references* = 105
  - *According to human evaluation of the titles and abstracts for each citation.

### Beta-Amyrin (CAS 559-70-6)

#### Compound names

Multiple names for beta-amyrin appear in the CAS Registry® (SciFinder®). These names are listed below along with the number of occurrences in the EndNote database (out of 1,854 total beta-amyrin citations):

- β-Amyrin
- Olean-12-en-3-ol, (3β)-
- Olean-12-en-3β-ol
- (+)-β-Amyrin
- 3β-Hydroxyolean-12-ene
- β-Amirin
- β-Amyrenol
- β-Amyrine

The names along with the number of occurrences in the EndNote database (out of 1,854 total beta-amyrin citations) are:

- Amyrin: 903 titles or abstracts
- Olean-12-en-3-ol: 2 titles or abstracts
- Hydroxyolean-12-ene: 6 titles or abstracts
- Amirin: 4 titles or abstracts
- Amyrenol: 1 title or abstract
- Amyrine: 9 titles or abstracts

The total occurrences of these compound names was 925 out of a total of 1,854 references. The remaining references may still be **OF-INTEREST** but lack the target name in the abstract or title.

#### SLM classifier phrases

The names, other than "amyrin" only amounted to 22 references (1%) of the total references. Since the SLM seemed to work better with fewer names, the following phrases were used for the small language model run:

- "Amyrin is present in plants."
- "Amyrin is not present in plants."

### Beta-Amyrone (CAS 638-97-1)

#### Compound names

Multiple names for alpha-amyrone appear in the CAS Registry® (SciFinder®). These names are listed below along with the number of occurrences in the EndNote database (out of 367 total beta-amyrone citations):

- β-Amyrone: 42 titles or abstracts
- Olean-12-en-3-one: 14 titles or abstracts
- 3-Oxoolean-12-ene: 2 titles or abstracts
- Pulcherrone: 0 titles or abstracts
- β-Amirenone: 1 titles or abstracts
- β-Amirone: 0 titles or abstracts
- β-Amyrenone: 43 titles or abstracts
- β-Amyron: 42 titles or abstracts

The total occurrences of these compound names was 144 out of a total of 367 references. The remaining references may still be **OF-INTEREST** but lack the target name in the abstract or title.

#### SLM classifier phrases

The following phrases were used for the small language model run:

- "Amyrone, olean-12-en-3-one, 2-oxoolean-12-ene, amirenone, amyrenone, or amyron is present in plants."
- "Amyrone, olean-12-en-3-one, 2-oxoolean-12-ene, amirenone, amyrenone, or amyron is not present in plants."

### Dammarenediol II (CAS 14351-29-2)

#### Compound names

Multiple names for Dammarene II appear in the CAS Registry® (SciFinder®). These names are listed below:

- Dammaranediol II
- Dammar-24-ene-3,20-dio, (3β)
- (3β)-Dammar-24-ene-3,20-diol
- Dammar-24-ene-3β,20-diol, (20S)-
- (+)-Dammarenediol II
- Dammar-24-ene-3β,20-diol
- 20*S*-Dammarenediol
- 20*S*-Dammarenediol II
- Dammarenediol

The names along with the number of occurrences in the EndNote database (out of 144 total Dammarenediol II citations):

- Dammarenediol: 59 titles or abstracts.
- [14351-29-2]: 1 abstract
- Dammar-24-ene-3,20-diol: 0 titles or abstracts. However, this name does appear in other fields of 19 EndNote references

Note: "Dammar" appeared in 87 titles or abstracts, but this truncated name would give positive responses to compounds that contained "dammar" but were not Dammarenediol II. Therefore "dammar" was not used as a classifier phrase for the SLM.

#### SLM classifier phrases

The following phrases were used for the small language model run:

- "Dammarenediol is present in plants."
- "Dammarenediol is not present in plants."

### Friedelin (CAS 559-74-0)

#### Compound names

Multiple names for Friedelin appear in the CAS Registry® (SciFinder®). These names are listed below:

- Friedelin
- 24,25,26-Trinoroleanan-3-one, 5,9,13-trimethyl-, (4β,5β,8α,9β,10α, 13α,14β)-
- (4β,5β,8α,9β,10α,13α,14β)-5,9,13-Trimethyl-24,25,26-trinoroleanan-3-one
- D:A-Friedooleanan-3-one
- (-) Friedelin
- 3(2H)-Picenone, eicosahydro-4,4a,6b,8a,11,11,12b,14a-octamethyl-, (4R,4aS,6aS,6bR,8aR,12aR,12bS,14aS,14bS)-
- 3(2H) Picenone, eicosahydro-4,4a,6b,8a,11,11,12b,14a-octamethyl-, [4R-(4α,4aα,6aβ,6bα,8aα,12aα,12bβ,14aα,14bβ)]-
- 3-Oxofriedelane
- Friedelan-3-one
- Friedelanone
- Friedeline

The names along with the number of occurrences in the EndNote database (out of 846 total Friedelin citations) are:

- Friedelin: 440 titles or abstracts
- Trinoroleanan-3-one: 0 titles or abstracts (but 78 references in "any field")
- Friedooleanan-3-one: 2 titles or abstracts
- Picenone: 0 titles or abstracts
- Friedelan-3-one: 43 titles or abstracts
- Friedelanone: 15 titles or abstracts
- Friedeline: 16 titles or abstracts

The total title/abstract citations with these names is 516 out of a total of 846 citations in the EndNote database. The remaining references may still be **OF-INTEREST** but lack the target name in the abstract or title.

#### SLM classifier phrases

The following phrases were used for the small language model run:

- "Friedelin, friedooleanan-3-one, friedelan-3-one, friedelanone, or friedeline is present in plants."
- "Friedelin, friedooleanan-3-one, friedelan-3-one, friedelanone, or friedeline is not present in plants."
