## Supplemental File 3 for "Small language models enable rapid and accurate extraction of structured data from unstructured text: an example with plants and their specialized metabolites"

***Evaluation of “maybe reports an occurrence” references using full texts***

Finally, to gain further insight into the classification process, the full text PDF files were acquired for the “maybe reports an occurrence” references for α-amyrone and dammarenediol II. Based on the content of the full-text PDF files, we resolved each of these “maybe reports an occurrence” references as either “reports an occurrence” or “does not report an occurrence” references. We elected to conduct this analysis for only two of case study triterpenoids since including the remaining two would have required downloading and manually examining more than 250 full-text articles, and more than 100 full-text articles were already being analyzed to assess α-amyrone and dammarenediol II “maybe reports an occurrence” references. For α-amyrone, 50 out of 70 ““maybe reports an occurrence”” references (71%) had plant/compound association data, while for dammarenediol II, 16 out of 34 “maybe reports an occurrence” references were found to contain plant/compound data (47%). Although in the case of α-amyrone several full text articles could not be retrieved, this analysis did suggest that a substantial portion of the number of articles initially flagged as “maybe reports an occurrence” that were in fact “reports an occurrence” references, based on full text inspection. Interestingly, many of the references that turned out to be “reports an occurrence” references were from projects that involved multiple plants and/or multiple compounds, and too much detailed information existed to include in the abstract. To build a database of compound species associations, these articles are, of course, of very high interest since they are rich in the type of information desired for such a database.

As a future direction, it may be possible to design a separate small language model-based workflow to classify “maybe reports an occurrence” references specifically, as part of a different filtering step. One possible approach to such an objective would be to use a higher number of classifier phrases, some of which are specifically designed to flag subtypes of articles for which the user wishes to specify a specific fate. This could potentially be used in combination with some type of machine learning classification model (a random forest, for example) trained on the output of the small language model that is assigning scores to a large number of classifier phrases. We briefly attempted this approach, but a challenge with such an approach is that abstracts, even if they describe similar projects, can be written in quite different ways. This diversity means that the ability of a small language model to recognize specific types of articles may be highly dependent on the way the abstract is written. This abstract variation in turn implies that a variety of filters would be needed to pick out specific article types of interest and that to automate the handling of references with middling scores in such a way would require a substantial and dedicated effort beyond the scope of the present work.
