## Supplementary figures and images for "Small language models enable rapid and accurate extraction of structured data from unstructured text: an example with plants and their specialized metabolites"

### Supplemental File 7

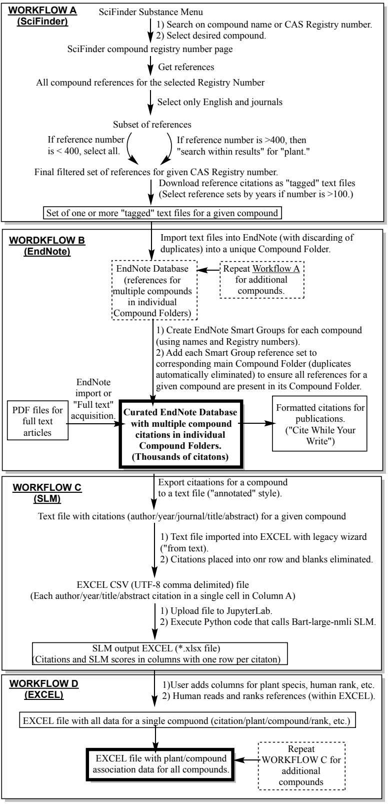
